# Integrating UHPLC-MS and MALDI-MSI for Spatial Nucleoside Profiling in FFPE Breast Cancer: A Multimodal Molecular Pathology Framework

**DOI:** 10.64898/2026.01.28.701949

**Authors:** Jierong Wu, Sabrina Y. Geisberger, Guido Mastrobuoni, Kamil Lisek, Sandra Raimundo, Grit Nebrich, Marta Grzeski, Nikolaus Rajewsky, Frederick Klauschen, Oliver Klein, Stefan Kempa

## Abstract

Formalin-fixed, paraffin-embedded (FFPE) tissues constitute the primary material for diagnostic pathology and retrospective clinical research, yet their use in metabolomics remains limited due to molecular cross-linking and analyte degradation. Here, we establish a cost-efficient molecular pathology workflow that integrates ultra-high-performance liquid chromatography mass spectrometry (UHPLC-MS) with matrix-assisted laser desorption/ionization mass spectrometry imaging (MALDI-MSI) to quantify and spatially map nucleosides in FFPE breast cancer tissues. Optimized extraction using methanol yielded nucleoside profiles comparable to fresh-frozen tissues, while MALDI-MSI enabled the spatial visualization of nine nucleosides across distinct histological regions. Several nucleosides including deoxyadenosine and 5-formylcytosine showed strong discriminatory power between tumor stages, revealing progressive metabolic rewiring during breast cancer progression.

Finally, spatial nucleoside patterns observed in a murine model were recapitulated in patient-derived FFPE tissues, underscoring the translational potential of nucleoside-based spatial metabolomics for clinical research and biomarker discovery. Together, this workflow establishes MALDI-MSI as a powerful and scalable spatial molecular pathology tool for interrogating nucleoside biology in archival breast cancer samples. Following MALDI-MSI, the same FFPE tissue sections can undergo laser capture microdissection, enabling genomic, proteomic, or targeted metabolomic profiling of MSI-defined tumor niches and microenvironmental regions. This integration directly links spatial nucleoside signatures to molecular alterations relevant to precision oncology in future.

## Introduction

Breast cancer remains one of the most common and heterogeneous malignancies affecting women around the world, with a diverse range of molecular subtypes and complex metabolic reprogramming.^1^ Nucleosides represent fundamental building blocks of nucleic acids and play a pivotal role in various cellular processes, including DNA replication, repair, and RNA transcription.^2^ The ability to directly detect nucleosides within tumor tissues has attracted increasing attention, providing new opportunities for metabolic characterization of cancer.

Recent studies have highlighted the emerging roles of nucleosides in cancer biology, where they participate in diverse processes including epigenetic modification, redox balance, and nucleotide salvage pathways, all of which are frequently dysregulated in tumors.^3,4^ Unmodified (canonical) nucleosides such as adenosine, cytidine, guanosine and uridine, as well as their deoxy forms, are central to primary nucleotide metabolism. They act as immediate substrates in nucleotide salvage pathways, enabling cells, especially rapidly proliferating tumor cells, to recycle nucleosides back into nucleotides efficiently.^5^ It is well known that cancer cells upregulate salvage pathways to meet the increased demand for DNA and RNA synthesis under nutrient-limited conditions. Pausing de novo synthesis and relying partly on salvage has been documented across multiple tumor types.^6^

Modified nucleosides, such as 5-methylcytidine and 5-formylcytosine in DNA and 5-methylcytidine and pseudouridine in RNA, are generated through enzymatic modification processes. They often reflect epigenetic or transcriptional regulation, rather than standard nucleotide metabolism.^7^ Unlike unmodified nucleosides, they are poor substrates for salvage pathways and are typically released or excreted during nucleic acid turnover. Their accumulation has been observed in cancer patients, and several studies have proposed them as potential non-invasive biomarkers for tumor detection and monitoring.^8^ As such, nucleosides have gained increasing attention not only as functional metabolites but also as promising biomarkers for cancer classification and staging.

In breast cancer research, nucleosides have been implicated in both therapeutic response and disease detection. Hsu et al. identified altered urinary nucleoside profiles in breast cancer patients, supporting their potential utility as non-invasive biomarkers.^9^ Similarly, Henneges et al. applied LC-MS-based urinary nucleoside profiling and identified statistically significant nucleoside signatures capable of distinguishing breast cancer patients from healthy individuals.^10^ However, the in situ composition and spatial distribution of nucleosides within breast cancer tissues remain poorly understood.

Indeed, the pronounced intertumoral heterogeneity of breast cancer, manifested in diverse histological patterns, molecular subtypes and metabolic phenotypes, presents a major challenge to accurate diagnosis and treatment stratification.^11^ In addition to such intertumoral variability, intra-tumoral heterogeneity, reflecting spatially distinct metabolic characteristics within individual tumors, often remains obscured in conventional metabolomic approaches based on bulk tissue.

Furthermore, the widespread use of FFPE in clinical research poses a significant challenge for the detection of small molecular metabolites due to extensive chemical cross-linking, molecular degradation, and reduced extraction of analytes.^12^ These challenges highlight the need for spatially resolved analytical strategies capable of preserving molecular context within FFPE tissues.

Among available spatially resolved techniques, matrix-assisted laser desorption/ionization mass spectrometry imaging (MALDI-MSI) offers a particularly effective solution for the direct, label-free visualization of metabolites within tissue sections.^13^ In contradistinction to techniques based on bulk analysis (such as UHPLC-MS) MALDI-MSI preserves spatial context, thereby enabling the simultaneous detection and localization of metabolites at cellular or even subcellular resolution.^14^ This technique has been successfully applied in various cancer studies to map metabolic heterogeneity, identify tumor margins, and explore drug distribution.^15,16,17^ As demonstrated in previous studies, MALDI-MSI has been employed to visualize spatial metabolomics and their applications in breast cancer, esophageal cancer and brain cancer. These studies have underscored the potential of MALDI-MSI in the field of spatial metabolomics.^18^ A significant advancement in this area was made by Alice Ly et al., who first established a robust protocol for the extraction and detection of metabolites from FFPE tissue. The spatial metabolomics workflow used in this study builds upon their approach.^19^

In the present study, we utilized complementary UHPLC-MS and MALDI-MSI analyses to investigate the distribution and metabolic dynamics of nucleosides in FFPE breast tumor tissues. The primary analysis was conducted on samples from a murine ductal carcinoma model^20^, with additional validation performed in a limited set of human breast cancer specimens. The combination of the high sensitivity and quantitative accuracy of UHPLC-MS with the spatial resolution of MALDI-MSI results in a comprehensive platform for nucleoside analysis in archival samples. This approach enables both precise quantification and spatial visualization of nucleosides, thereby offering new opportunities to explore tumor-specific metabolic reprogramming. Notably, previous attempts to profile primary metabolites using GC-MS produced limited results in FFPE tissues, emphasising the technical difficulties involved in detecting small molecules in preserved samples. By contrast, nucleosides produced stable and reproducible signals, rendering them better suited to spatial metabolomic analysis in this context. Taken together, our findings contribute to a deeper mechanistic understanding of breast cancer progression and support the translational potential of spatial nucleoside metabolomics in cancer research.

## Materials and Methods

### Sample Collection

Murine breast tumor tissues were kindly provided by the group of Prof. Nikolaus Rajewsky, PhD (Max Delbrück Center, Berlin Institute for Medical Systems Biology). The model was generated and characterized by K.L.,.^20^ Animal experiments were conducted in compliance with institutional and governmental guidelines. All procedures involving animals were approved by the Landesamt für Gesundheit und Soziales (LaGeSo), Berlin (approval no. G#0213.18). K.L. received financial support from the Deutsche Krebshilfe (grant no. 70113738). Tissues were derived from a transgenic mouse model conditionally activating Pik3caH1047R, Trp53R172H, and stabilized β-catenin alleles in the mammary epithelium under WAPiCre control, which recapitulates key features of triple-negative breast cancer (TNBC) progression, including ductal transformation and intra-tumoral heterogeneity.^20^

Tumors were preserved using two different methods: (1) formalin fixation followed by paraffin embedding (FFPE), and (2) embedding in Tissue-Tek O.C.T. compound and storage at −80 °C as fresh frozen tissues.^20^ UHPLC-MS was performed on both FFPE and fresh-frozen tissues, while MALDI-MSI was conducted exclusively on FFPE samples to assess nucleoside profiles and spatial distribution.

FFPE breast cancer samples from four patients (three TNBC and one luminal subtype) were obtained from the group of Prof. Frederick Klauschen, MD (Institute of Pathology, Ludwig Maximilian University of Munich). These samples were collected under appropriate ethical approvals and used exclusively for MALDI-MSI analysis to assess the detectability and spatial localization of nucleosides in clinical FFPE tissues. The study was approved by the Ethics Committee of LMU Munich (approval no. 23-0650).

### Nucleoside Extraction

FFPE breast tissue sections (10 µm thick) were prepared using a microtome (Leica, Germany) and placed into 1.5 mL microcentrifuge tubes. Nucleosides were extracted following the protocol described by Cacciatore S et, al.^21^ To assess the impact of deparaffinization, some samples were pretreated with xylene before extraction.

For extraction, 1 mL of 80% methanol was added directly to the samples, followed by incubation at 70°C for 40 minutes. The samples were then placed on ice for 15 minutes and centrifuged at 14,000g for 10 minutes at 4°C. The supernatant was transferred to a new 1.5 mL microcentrifuge tube and put on ice for 10 minutes, then underwent centrifugation at 14,000g for 5 minutes at 4°C. The final supernatant was collected, dried overnight, and stored at -80°C for further analysis (SFigure 1A).

Fresh frozen breast tissue samples were sectioned at 10 µm thickness using a CryoStar NX70 cryo-microtome (Thermo, USA) and placed into 1.5 mL microcentrifuge tubes. Nucleosides were extracted using 80% methanol, followed by incubation at room temperature for 4 hours. The samples were then centrifuged at 14,000g for 10 minutes. The supernatant was collected, dried overnight, and stored at -80°C for further analysis (SFigure 1B).

### Nucleoside Stock Solution

Stock solutions of individual ribonucleosides and deoxyribonucleosides (1 mg/mL each in water) were prepared and stored at –80 °C. For calibration and method evaluation, a mixed standard solution containing 30 nucleosides (19 modified ribonucleosides, 5 unmodified ribonucleosides, and 6 deoxyribonucleosides) was freshly prepared by combining the individual stocks and diluting them with 0.1% formic acid in water to yield a final concentration of 500 pg/µL per compound. Serial dilutions of this mixture were used to assess linearity, sensitivity, and reproducibility in subsequent UHPLC-MS analyses.

Samples were resuspended in 35µL 0.1% formic acid and then measured by UHPLC-MS.

### UHPLC-MS Analysis

Samples were measured with a Vanquish Horizon UHPLC system (Thermo Scientific, USA) coupled to a TSQ Quantiva triple quadrupole mass spectrometer (Thermo Scientific, USA).

Chromatographic separation was achieved with a Kinetex Biphenyl Column (2.6 μm, 100 × 2.1 mm, Phenomenex). The mobile phase consisted of 0.1% acetic acid in water (A) and 0.1% acetic acid in methanol (B), and a linear gradient was applied as follows: from 0% to 30% B over 4.5 minutes, 30% to 50% B in 0.5 minutes, held at 50% B for 0.5 minutes, and returned to 0% B for 2 minutes of re-equilibration. The flow rate was set to 300 μL/min, and the column temperature was maintained at 30°C with a sample injection volume of 1 μL.

The mass spectrometer was operated in both positive and negative ionization modes using a heated electrospray ionization (HESI) source with the following optimized settings: 3500 V (positive mode) and -2500 V (negative mode) for the ionization voltages, with sheath gas, auxiliary gas, and sweep gas at 30 Arb, 10 Arb, and 1 Arb, respectively. The ion transfer tube temperature was set to 325°C, the nebulizer gas was nitrogen, and the collision gas was argon (1.5 mTor) at a vaporizer temperature of 350°C. Nucleosides were quantified using selected reaction monitoring (SRM) mode with optimized parameters (precursor m/z, product m/z, and collision energy for each nucleoside) as listed in Supplementary Table.

### Histology and Immunostaining

Mammary glands were fixed overnight at 4°C in 4% formaldehyde (Roth), then dehydrated and embedded in paraffin. For staining, tissue sections were deparaffinized, rehydrated, stained with Hematoxylin and Eosin (H&E), dehydrated, and mounted. Images were acquired with a bright-field Zeiss microscope. Cryosections were directly frozen in OCT after tissue was excised from the mice. Before the staining sections were fixed with 4% methanol-free paraformaldehyde for 15 minutes. All samples were blocked with 10% horse serum for 1 hour at room temperature before overnight incubation with primary antibodies at 4°C. Antibodies: mouse E-cadherin/Cdh1, rabbit MMP9, goat GFP (that recognizes YFP protein) were used in 1:50 dilution. For fluorescent staining, samples were incubated with secondary antibodies and DAPI for 1 hour at room temperature before mounting with Immu-Mount (Thermo Scientific).

### Immunofluorescence imaging

Immunofluorescence imaging Fluorescently stained cryosections were imaged with high-resolution confocal imaging using a Leica Stellaris 5 system equipped with a White Light Laser and HyD detectors. Sections were scanned using a 20× HC PL APO CS2 glycerol objective (NA 0.75) with 1024×1024 pixel resolution and sequential laser scanning. Images were acquired at 20× magnification and stitched using Leica LAS X software when required for whole-gland visualization. Z-stacks were collected when necessary for visualization of spatial organization within tumor-stromal niches. All image acquisitions were performed using standardized laser settings and exposure times to allow quantitative comparisons across conditions.

### MALDI-MSI Analysis

Samples were prepared as previously described Alice Ly et,al.19 FFPE blocks were sectioned at a thickness of 4 μm using a microtome. Sections were mounted onto indium-tin-oxide (ITO) slides (Bruker Daltonik, Bremen, Germany) and put in a 40 °C oven over night. Mounted slides were dried in a 60 °C oven for 1 hour to promote tissue adhesion. Deparaffinization was performed by immersing the slides in fresh xylene for 8 minutes, followed by a second 8-minute incubation in a separate xylene bath. Slides were then put on a 40 °C hot plate to dry prior to further processing. To aid with peak identification, a droplet of nucleoside standard mixture was spotted onto the edge of each ITO slide, outside the tissue section (SFigure 1C).

Matrix solution (10 mg/mL α-cyano-4-hydroxycinnamic acid in 70% acetonitrile, 0.1% trifluoroacetic acid, heated to 60 °C) was applied to tissue sections using an HTX TM-Sprayer (HTX Technologies) under uniform conditions. During matrix deposition, a recrystallization chamber was prepared using a large plastic Petri dish lined with a round paper filter and equipped with four small magnets. After spraying, slides were subjected to recrystallization in 5% isopropanol vapor to enhance matrix crystal homogeneity and analyte co-crystallization.

After matrix application, ITO slides were fixed to the lid of a plastic Petri dish using small magnets, ensuring the tissue side faced downward. A round filter paper pre-wetted with 5% isopropanol was placed inside the base of the dish to generate vapor. The closed chamber was then incubated at 55 °C for 1 minute and 15 seconds to allow recrystallization of the matrix.

Analyses were performed using a RapifleX MALDI Tissuetyper (Bruker Daltonik GmbH, Bremen, Germany) equipped with flexImaging 5.1 and flexControl 3.0 software in positive ion reflector mode with a detection range of 600–1200 m/z, 500 laser shots per point, a sampling rate of 1.25 GS/s (gigasamples per second), and a raster width of 50 µm for MALDI-MSI data acquisition.

External calibration was performed using phosphor red standards (Bruker Daltonik GmbH). After matrix removal, Sections were stained with hematoxylin and eosin (H&E) for histological examination.

### Data Processing

UHPLC-MS raw data were processed using Thermo Xcalibur Quan Browser, following the Quantitative Analysis User Guide provided by the manufacturer. Briefly, extracted ion chromatograms (XICs) were generated targeting each nucleoside with a mass tolerance of ±5 ppm. Peak integration parameters, such as baseline subtraction, smoothing algorithm, and peak boundaries, were manually reviewed and adjusted using the integrated peak detection settings. Calibration curves constructed from authentic standards and internal standards enabled absolute quantification of nucleoside concentrations. Peak area data were exported and visualized using custom R scripts for downstream statistical analysis.

MALDI-IMS raw data were imported into SCiLS Lab Pro (version 2025b, Bruker Daltonik GmbH) with total ion count (TIC) normalization enabled and baseline removal disabled. The data were converted to .sbd and .slx formats for further processing. Spectral alignment was performed across the dataset using a bin width of 0.05 Da to group nearby peaks. Manual peak picking was then conducted based on aligned spectral features.

To identify discriminatory m/z features between tumor and normal tissue regions, supervised ROC analysis was applied to the partitioned datasets. To ensure balanced group comparison, 800 m/z values were randomly selected per region, based on the size of the smallest group. Features with an AUC > 0.7 or < 0.3 were further evaluated using a Wilcoxon rank-sum test to assess statistical significance. Alignment of MALDI-MSI and UHPLC-MS m/z values was accepted with a mass difference ≤0.3 Da. Isobaric nucleosides with overlapping m/z values could not be resolved by MALDI-MSI and were therefore reported together throughout the analysis.

## Results

### Effect of Extraction Methods on Nucleoside Detection

We evaluated the effect of two different experimental methods on nucleoside extraction (methanol (MeOH) extraction and conventional xylene deparaffinization) in FFPE breast cancer tissues and using fresh frozen tissue extraction with methanol as a control group.

As shown in Figure 1A, the overall nucleoside signal intensity was lower in FFPE tissues compared to fresh-frozen samples, independently of the extraction method. Only minimal differences were observed between the two extraction approaches, with methanol extraction yielding slightly higher signal intensities and a metabolic profile more comparable to that of fresh-frozen tissues. Based on these results, methanol extraction was selected for UHPLC-MS analysis.

**Figure 1.**
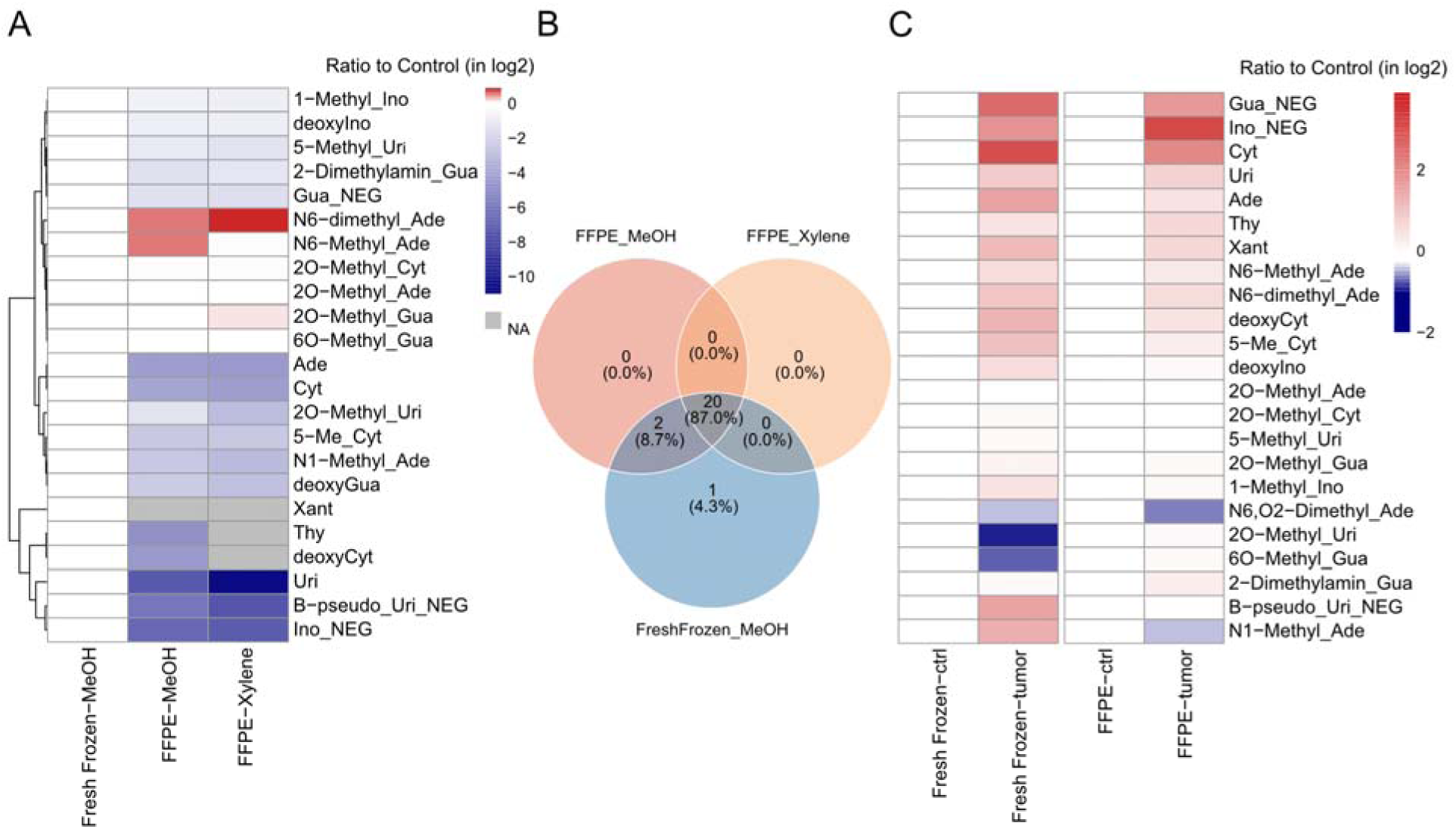
Comparison of extraction protocols and tissue preservation on nucleoside profiling. (A) Heatmap showing log□-transformed nucleoside intensities in FFPE tissues processed via methanol (FFPE_MeOH), xylene (FFPE_Xylene), and fresh frozen samples (Fresh Frozen_MeOH). (B) Venn diagram of shared nucleosides detected across the three groups. (C) Heatmap comparing tumor and normal tissues across FFPE and fresh-frozen samples, shown as log□-transformed ratios relative to control (normal) tissues.

For MALDI-MSI, however, xylene-based deparaffinization was applied following the established protocol (Ly et al.)^19^ to ensure tissue morphology preservation and compatibility with matrix application. Attempts to perform MALDI analysis on non-deparaffinized FFPE sections resulted in a complete loss of signal, confirming the necessity of paraffin removal for successful ionization. Although the two preparation strategies are not fully equivalent, this combination provided a practical and consistent workflow for nucleoside analysis in FFPE specimens.

We compare nucleosides found in fresh frozen tissue, FFPE tissues extracted with methanol (FFPE_MeOH), and FFPE tissues processed by xylene deparaffinization (FFPE_Xylene) (Figure 1B). A total of 20 nucleosides were identified in all three groups concurrently. Furthermore, a mere two nucleosides were detected in fresh frozen and FFPE_MeOH, while no overlap was observed between fresh frozen and FFPE_Xylene alone. It is note that the presence of a nucleoside in the fresh frozen group suggests that some specific substances may have been lost or changed during the preparation of FFPE samples.

Next, we compared the relative change of nucleosides in tumor tissues compared to normal tissues in FFPE and fresh frozen samples (Figure 1C). The most nucleosides show an upregulated trend in tumor tissues. The ratios of nucleosides in FFPE samples are generally similar to those in fresh frozen tissue. The results indicate that key metabolic signatures which distinguish between tumor and normal tissues are largely retained in FFPE tissues when extraction strategies have been optimized.

These findings suggest that methanol extraction slightly improves the extraction of nucleosides from FFPE samples. Although FFPE preservation allows for analysis of previously stored tissues, it can also affect the detection of certain unstable or modified nucleoside types.

### Spatial Distribution of Nucleosides in FFPE Mouse Samples

In order to investigate the spatial distribution of nucleosides in breast cancer tissues, we performed MALDI-MSI on FFPE tissue sections. 9 nucleosides were detected by MALDI-MSI, with peak assignments initially based on matching to theoretical m/z values and subsequently confirmed by UHPLC-MS. The observed mass deviations varied across compounds, ranging from as low as 0.078 Da to a maximum of 0.2 Da, reflecting differences in ionization behavior, local tissue environment, and the distance from the external reference spot. Some nucleosides from the spotted standard mix were not detected, likely due to matrix effects or surface ion suppression. Thus, peak identity was further confirmed by UHPLC-MS and spatial consistency in tissue sections. All detected nucleosides were further supported by independent UHPLC-MS confirmation. The analysis of ion images has revealed different spatial distribution patterns across the tissue, with several nucleosides concentrated in areas of high tumor density (SFigure 2). The findings suggest that MALDI-MSI provides high-resolution spatial information on nucleoside metabolism in FFPE tissues, capturing not only tumor-associated enrichment but also intratumor heterogeneity.

Next, nucleoside localization and abundance were compared at different pathological stages of ductal carcinoma, including normal breast tissue, early, intermediate, and advanced tumors (Figure 2). Representative MSI ion images were aligned with H&E-stained sections in order to delineate tumor boundaries and highlight histological features.

**Figure 2.**
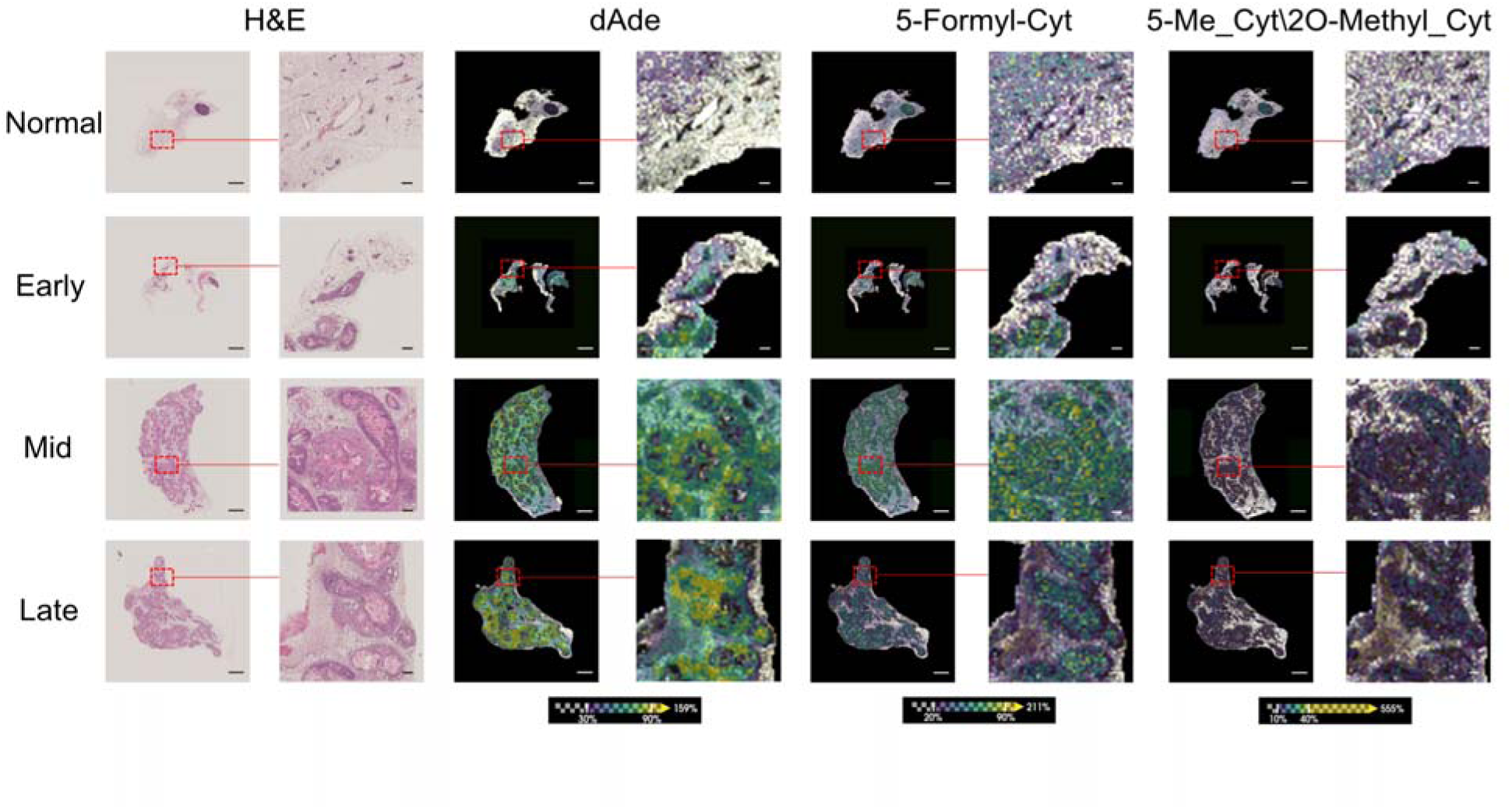
Comparative analysis of nucleoside levels across tumor progression stages using MALDI-MSI. Representative tumor regions were selected from MALDI-MSI datasets to compare the relative intensities of deoxyadenosine (dAde), 5-formylcytosine and 5-methylcytidine\2O-methylcytidine across early, mid, and late stages of breast cancer. Ion images illustrate stage-specific changes in nucleoside abundance. Scale bars: 2 mm (overview images) and 200 µm (magnified views).

A progressive increase in nucleoside spatial intensity was observed across tumor stages, with minimal signals in normal breast ducts, modest elevations in early-stage tumors, and marked accumulation in viable regions of intermediate and late-stage tumors. This pattern was consistently detected in several nucleosides, including deoxyadenosine, 5-formylcytosine, and 5-methyl_cytidine\2O-methyl_cytidine, suggesting a potential link between nucleoside accumulation and metabolic reprogramming during tumor progression.

This suggests that using MALDI-MSI with a histopathological context can visualize metabolic changes that depend on the stage of the disease, providing valuable insights into how breast cancer progresses.

### Diagnostic Potential of Nucleosides for Stratifying Breast Cancer Progression

We performed receiver operating characteristic (ROC) curve analysis based on MALDI-MSI-derived ionic strength to assess the diagnostic relevance of nucleosides at different stages of breast cancer (SFigure 3). The ROC curve is utilized to evaluate the capacity of a marker to differentiate between distinct pathological states by plotting the relationship between sensitivity and (1−specificity). The area under the curve (AUC) can be employed to quantitatively assess its classification efficacy, and AUC value greater than 0.7 was indicative of high expression in tumors.

In this study, the ROC curves were plotted and statistically tested. We compared nucleoside distribution differences between normal breast tissue and early, intermediate, and advanced breast ductal carcinoma tissues, then further compared differences between early, middle, and advanced tumors to identify nucleosides with stage specificity.

Among the nine nucleosides detected by MALDI-MSI, several exhibited strong discriminatory power for distinguishing different tumor stages. Notably, 5-formylcytosine and deoxyadenosine achieved AUC values of 0.705 and 0.715, respectively, when comparing early-stage tumors to normal tissue, indicating their potential as early diagnostic markers (Figure 3A). In addition, 5-methylcytidine\2O-methylcytidine showed an AUC of 0.702 in differentiating late-stage tumors from early-stage ones (Figure 3B), while deoxyadenosine yielded an AUC of 0.267 when comparing mid-stage to early-stage tumors (Figure 3C), suggesting a possible decrease during intermediate progression.

**Figure 3.**
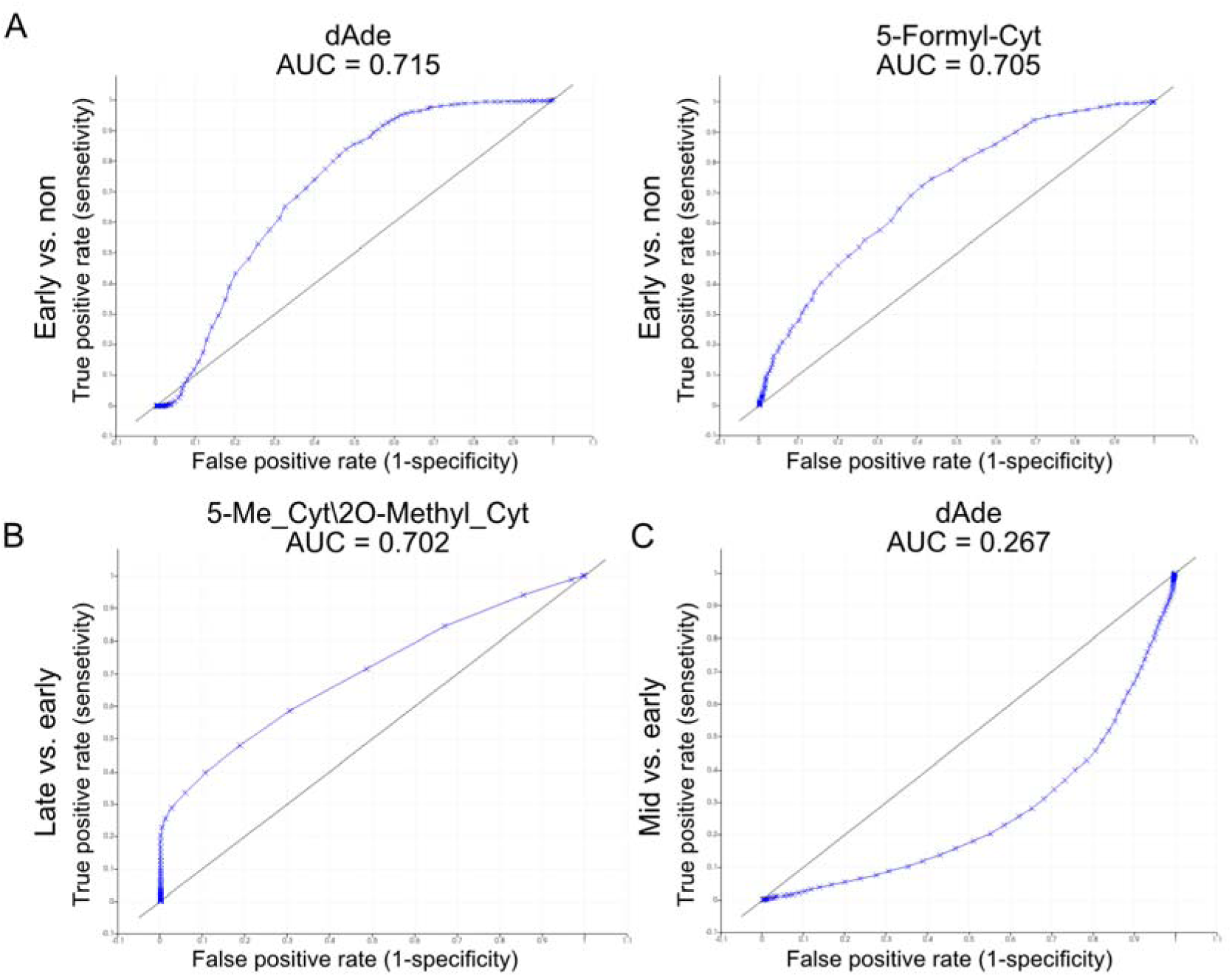
Receiver operating characteristic (ROC) curves of selected nucleosides for tumor stage classification. (A) ROC curves for deoxyadenosine (dAde) and 5-formylcytosine in distinguishing early-stage tumors from normal tissue, showing AUC values of 0.715 and 0.705, respectively. (B) ROC curve for 5-methylcytidine\2O-methycytidine (5-Me_Cyt\2O-Methyl_Cyt) in differentiating late-stage tumors from early-stage tumors, with an AUC of 0.702. (C) ROC curve for deoxyadenosine (dAde) in distinguishing mid-stage tumors from early-stage tumors, with an AUC of 0.267, indicating potential value in stratifying tumor progression. Each curve illustrates the trade-off between sensitivity and specificity across threshold values. AUC values closer to 1 or 0 indicate higher discriminative power.

Taken together, a subset of the nine nucleosides demonstrated notable stage-associated changes, underscoring their potential utility as auxiliary biomarkers for tumor diagnosis or progression assessment. To visualize their stage-specific abundance more intuitively, we further generated box plots illustrating the spatial intensity distributions across different tumor stages (SFigure 4).

Several nucleosides displayed a progressive increase in abundance as tumor stage advanced. For instance, 5-formylcytosine showed significantly elevated levels in early tumors compared to normal tissue (p < 0.01). This observation aligns with previous reports describing 5-formylcytosine as an intermediate of active DNA demethylation involved in epigenetic regulation.^22^ Similarly, xanthosine and inosine were markedly upregulated in mid and late-stage tumors, consistent with previous reports linking these nucleosides to enhanced purine metabolism in proliferating cells.^23^ Other nucleosides, such as 5-methylcytidine and deoxyadenosine, also followed similar trends, further supporting the relevance of nucleoside dynamics in malignancy development. These findings provide further evidence that tumor stage-specific metabolic rewiring can be captured via spatial nucleoside profiling.

In conclusion, the observed nucleoside signatures reflect dynamic changes during breast cancer progression, and may offer practical value as stage-related markers or auxiliary molecular indicators for spatial metabolomic profiling.

### Spatial Validation of Nucleoside Distributions in Human FFPE Breast Cancer Samples

To evaluate the translational relevance of our findings, we extended MALDI-MSI analysis to five FFPE human breast cancer tissue samples from clinical patients, including four triple negative breast cancer (TNBC) and one luminal subtype. Among the three nucleosides that demonstrated strong stage discrimination in mouse ROC analysis, deoxyadenosine and 5-formylcytosine were also identifiable in patient FFPE samples, exhibiting distinct spatial localization patterns within tumor regions. The alignment with corresponding H&E-stained sections allowed for clear localization of nucleoside signals relative to histopathological features.

Spatial ion images revealed marked heterogeneity in nucleoside distribution across patients (SFigure 5). Within the same molecular subtype (TNBC), variations in signal intensity and localization were observed, suggesting substantial inter-patient metabolic diversity. Notably, certain nucleosides such as deoxyadenosine and 5-formylcytosine were enriched in tumor regions across most TNBC samples, whereas their spatial signals in the luminal case appeared more diffuse or restricted (Figure 4). These findings indicate that nucleoside metabolism in breast cancer may be modulated not only by tumor progression but also by molecular subtype and individual tumor microenvironment.

**Figure 4.**
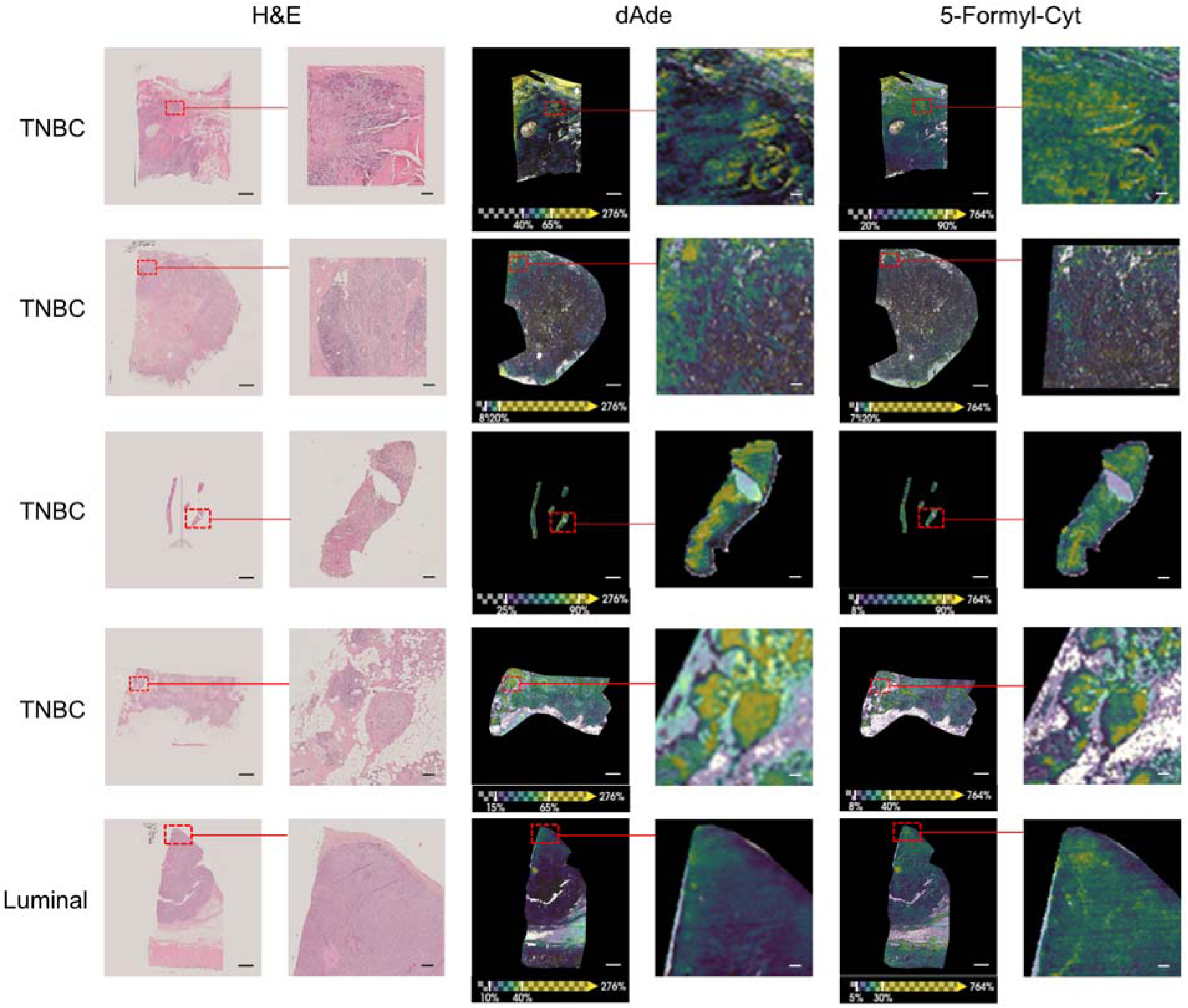
Spatial distribution of nucleosides in clinical breast cancer tissues analyzed by MALDI-MSI. Representative tumor regions from five FFPE breast cancer patients (four TNBC and one luminal subtype) were selected to evaluate spatial patterns of dAde and 5-formyl-cytosine by MALDI-MSI. Enlarged views show nucleoside enrichment in viable tumor areas. Scale bars: 2 mm (overview images) and 200 µm (magnified views)

All ion images were normalized to TIC to ensure comparability across sections. The representative examples of deoxyadenosine and 5-formylcytosine highlight that key nucleoside patterns observed in animal models can be recapitulated in clinical samples, supporting their potential as stage- or subtype-associated metabolic markers.

## Discussion

In this study, we employed a combined UHPLC-MS and MALDI-MSI platform to achieve quantitative analysis and spatial mapping of nucleosides in FFPE breast cancer tissues. While UHPLC-MS offers high sensitivity and specificity for nucleoside quantification, MALDI-MSI complements this by preserving tissue architecture and enabling metabolite localization with high spatial resolution, providing a more comprehensive metabolic landscape.^24^ While UHPLC-MS was performed in both positive and negative ionization modes to ensure comprehensive detection, MALDI-MSI acquisition was limited to positive ionization mode. This decision was based on the ionization behavior of nucleosides, the majority of which are preferentially ionized in positive mode, whereas only a small subset show detectable signals in negative mode.

Focusing on positive mode thus allowed us to target the dominant nucleoside species with higher sensitivity and efficiency. In addition, the use of α-cyano-4-hydroxycinnamic acid (CHCA) as matrix, known for its vacuum stability and homogeneous crystallization, further supports consistent positive-mode acquisition across large tissue cohorts. This dual approach builds upon recent advances in spatial metabolomics, which have demonstrated the power of combining mass spectrometric imaging with targeted metabolite quantitation to unravel tumor heterogeneity and metabolic rewiring.^25^

Optimizing extraction protocols was critical for maximizing nucleoside recovery from FFPE tissues, which typically pose challenges due to cross-linking and formalin-induced modifications.^26^ Our findings indicate that methanol extraction produced stronger and more consistent nucleoside signals compared to conventional xylene-based deparaffinization, consistent with previous observations that alcohol-based protocols can improve metabolite recovery and preservation in FFPE specimens.^27^ Importantly, the nucleoside profiles obtained from FFPE tissues using optimized methods closely resembled those from fresh-frozen samples, reinforcing the reliability of FFPE samples for metabolomic analysis. Methanol extraction was applied for UHPLC-MS to maximize nucleoside recovery, whereas xylene-based deparaffinization was used for MALDI-MSI to maintain tissue integrity and matrix compatibility. Although the two approaches differ, this combination provided a practical balance between analytical performance and sample preservation in FFPE nucleoside analysis.

Spatial mapping by MALDI-MSI revealed heterogeneous distribution of nucleosides in breast tumor sections, with enhanced signals predominantly localized to the tumor periphery. These patterns likely reflect metabolic gradients arising from differential oxygenation, nutrient availability, and cell viability within tumors, as supported by prior studies demonstrating metabolic zonation in solid malignancies.^28^ Interestingly, while some nucleosides showed strong signal intensities throughout the tumor mass, a marked decrease in central regions was frequently observed. These areas are presumed to be necrotic, a common feature in large solid tumors, which often exhibit reduced metabolic activity and disrupted tissue architecture. In contrast, nucleoside accumulation in viable peripheral regions likely reflects higher metabolic turnover or increased preservation of nucleic acid metabolites at the proliferative tumor margin.

To further contextualize these spatial patterns, we compared MALDI-MSI nucleoside signals with multiplex immunofluorescence staining performed on tumor sections from the same transgenic model (SFigure 6). Although the MALDI-MSI and immunofluorescence data were not acquired from the identical tissue section, both modalities consistently demonstrated peripherally enriched regions that correspond to viable, epithelial-like tumor areas with high E-cadherin and YFP signals, and inner regions with reduced marker intensity that are compatible with necrotic or stromal composition. This orthogonal comparison supports the biological relevance of the spatial ion maps and illustrates the feasibility of integrating metabolite imaging with phenotypic markers. Although exploratory, this multimodal approach opens possibilities for downstream multi-omic profiling, such as combining MALDI-MSI with laser capture microdissection (LCM), proteomics, or spatial transcriptomics in FFPE samples.

Such spatial differences underscore the value of MALDI-MSI in resolving intra-tumoral metabolic heterogeneity; a feature that is lost in bulk metabolomic profiling space. Decreased nucleoside intensity in central tumor regions is consistent with necrotic tissue characteristics, highlighting the ability of MALDI-MSI to resolve spatially distinct metabolic states within tumors.^29,30^

Several nucleosides were found to be progressively upregulated during tumor development.^31^ 5-formylcytosine, an intermediate in TET-mediated active DNA demethylation, was found to be elevated at early tumor stages, suggesting early epigenetic remodeling during breast cancer development.^32,33^ This finding is consistent with evidence that 5-formylcytosine may accumulate in specific tumor contexts and influence genome structure and gene regulation through its role as an epigenetic modification.^34,35^ As a modified nucleoside, 5-formylcytosine reflects changes in RNA turnover rather than canonical nucleotide metabolism. In contrast, while deoxyadenosine (dAde) has not previously been widely reported as a cancer-stage biomarker, its metabolic role as a purine nucleoside and its dynamic regulation during cell proliferation suggest that alterations in deoxyadenosine levels may reflect underlying shifts in nucleotide metabolism during tumor progression.^36^

A subsequent diagnostic evaluation incorporating ROC analysis further underscored the potential of spatial nucleoside profiling in tumor staging. 5-formylcytosine and deoxyadenosine exhibited high discriminative power between normal and early-stage tumors, with AUC values exceeding 0.7. In addition, 5-methylcytidine/2O-methylcytidine was preferentially elevated in advanced-stage tumors, indicating potential involvement in later-stage tumor biology, possibly linked to RNA methylation processes or aberrant nucleic acid turnover.^37,38^ These findings are consistent with prior reports proposing modified nucleosides as promising metabolic indicators for tumor progression and patient stratification.^39^ By integrating MALDI-MSI with quantitative mass spectrometry, we achieved comprehensive profiling of nucleoside abundance while preserving spatial heterogeneity; an increasingly recognized factor in tumor progression and therapeutic resistance.^40,41^

In addition to murine models, spatial nucleoside profiling was also applied to clinical FFPE breast cancer samples. Although no statistical analysis was performed due to limited sample size and lack of matched controls, the results nonetheless demonstrated consistent detection of several nucleosides, including deoxyadenosine and 5-formylcytosine, across patient tumors. It is noteworthy that spatial ion images revealed considerable inter-patient heterogeneity in both signal intensity and localization, which may be indicative of differences in tumor subtypes, metabolic background, or microenvironmental factors. These observations underscore the complexity of nucleoside metabolism in clinical contexts and highlight the feasibility of extending spatial metabolomics workflows to patient-derived FFPE specimens.

Finally, the successful application of MALDI-MSI to FFPE sections highlights the expanding role of spatial metabolomics in translational oncology. While fresh-frozen tissues remain the gold standard for molecular imaging, the ability to extract meaningful metabolic information from FFPE archives opens new avenues for retrospective studies correlating molecular profiles with long-term clinical outcomes.^42,43^ As spatial resolution, mass accuracy, and analytical pipelines continue to improve, integrated spatial metabolomics strategies are poised to become a cornerstone in cancer research and biomarker discovery.

Beyond the biological insights gained in this study, an important strength of our workflow lies in its methodological scalability and compatibility with multimodal downstream analyses. MALDI-MSI represents a relatively cost-efficient spatial omics technology compared with alternative high-resolution imaging platforms, requiring minimal consumables and enabling high-throughput measurement of large FFPE tissue cohorts.

Once MSI acquisition is complete, the tissue section remains fully accessible for additional analyses, positioning MALDI-MSI as a powerful entry point into serial multimodal profiling of the same specimen. In particular, the compatibility of MALDI-MSI with laser capture microdissection (LCM) enables precise isolation of histologically or metabolically defined regions, such as invasive fronts, hypoxic niches, or stromal microenvironments, identified directly from the ion images. These micro-dissected regions can then be subjected to proteomics, metabolomics, or targeted genomic analyses, including next-generation sequencing, methylation profiling, or precision oncology panels, thereby linking spatial nucleoside features with tumor-specific genetic and transcriptional programs.

This integrative framework is especially valuable for FFPE material, where the ability to reuse the same section maximizes information gain from limited archival tissue while directly connecting MSI-derived metabolic phenotypes to genomic drivers of tumor heterogeneity and microenvironmental remodeling. By enabling spatially resolved nucleoside imaging that can be seamlessly followed by high-content molecular assays, our approach provides a flexible and translationally relevant strategy for dissecting tumor ecosystems and identifying clinically actionable metabolic–genomic interactions in breast cancer.

## Conclusion

In summary, the present study substantiates the viability of employing an integrated UHPLC-MS and MALDI-MSI approach to profile and spatially map nucleoside distributions in FFPE breast cancer tissues. The findings of this study demonstrate the presence of dynamic metabolic alterations in nucleosides across various stages of tumor progression in a murine model.

Furthermore, the study substantiates the reliability of detecting key nucleosides, such as deoxyadenosine and 5-formylcytosine, in clinical FFPE samples. The observed spatial heterogeneity and inter-patient variation highlight the complexity of tumor metabolism and underscore the translational relevance of nucleoside-based spatial metabolomics. The capacity to procure both quantitative and spatially resolved information unveils novel avenues for cancer metabolomics research and furnishes a robust platform for investigating metabolic rewiring in tumors. Future studies should aim to validate these findings in larger and more diverse patient cohorts, and to further explore the diagnostic and prognostic potential of nucleoside-based biomarkers in breast cancer.

## Supporting information

SFigure1

SFigure2

sFigure3

SFigure4

SFigure5

SFigure6

STable1

## ASSOCIATED CONTENT

The following files are available free of charge.

**SFigure1. Sample preparation and analysis workflow.**

(A) Workflow for nucleoside extraction from FFPE tissue sections (10 µm), with or without xylene-based deparaffinization.

(B) Workflow for fresh-frozen tissue extraction.

(C) FFPE tissue sections (4 µm) on ITO-coated slides underwent deparaffinization, matrix application, and MALDI-MSI analysis, followed by H&E staining and data processing for spatial metabolite distribution analysis. (file type, PDF)

**SFigure 2. Spatial distribution of nucleosides in breast tissue samples at different tumor stages.**

Representative MALDI-MSI ion images of 9 nucleosides detected in FFPE breast tissues across normal, early, mid, and late tumor stages. Corresponding H&E-stained images are shown for histological reference. Scale bars: 2 mm (overview) and 200 µm (magnified). (file type, PDF)

**SFigure 3. ROC analysis of nucleoside signals for tumor stage classification.**

Receiver operating characteristic (ROC) curves for individual nucleosides based on MALDI-MSI intensity data. The area under the curve (AUC) is indicated for each comparison. (A) Early-stage tumors vs. normal tissue; (B) Mid-stage tumors vs. normal tissue; (C) Late-stage tumors vs. normal tissue; (D) Late-stage tumors vs. early-stage tumors; (E) Late-stage tumors vs. mid-stage tumors; (F) Mid-stage tumors vs. early-stage tumors.

Nucleosides with AUC values greater than 0.7 or less than 0.3 were considered to have potential as stage-specific biomarkers. (file type, PDF)

**SFigure 4. Box plots illustrating the normalized signal intensities of nucleosides detected by MALDI-MSI across different stages of tumor progression.**

Samples were categorized into normal, early-stage tumor, mid-stage tumor, and late-stage tumor groups. Each plot displays the distribution of nucleoside intensities across tissue regions, with blue points representing individual pixel measurements and red points denoting statistical outliers. Statistical comparisons between groups were performed using unpaired two-sided t-tests. Significance levels are indicated as follows: p < 0.05 (*), p < 0.01 (**), p < 0.001 (***), and ns = not significant. (file type, PDF)

**SFigure 5. Overview of spatial distribution of nucleosides in FFPE clinical breast cancer tissues.**

MALDI-MSI ion images of 10 nucleosides detected in FFPE tissue sections from five breast cancer patients (four TNBC and one luminal subtype). All ion images were normalized to total ion current (TIC) and displayed on a common intensity scale. Scale bars: 2 mm (overview) and 200 µm (magnified). (file type, PDF)

**SFigure 6. Multimodal visualization of deoxycytidine distribution and phenotypic tumor markers in FFPE murine breast tumors.**

(A) MALDI-MSI ion image showing the spatial distribution of deoxycytidine (dCyt) in a representative FFPE breast tumor section from a transgenic murine model, together with the corresponding H&E stain of the same section after matrix removal. Scale bars: 400 µm.

(B) Multiplex immunofluorescence image from a tumor section obtained from a different mouse of the same transgenic model, stained for E-cadherin (red), MMP9 (green), and YFP (yellow), with individual channels shown on the right. These markers highlight epithelial structures (E-cadherin□), proteolytically active tumor/stromal regions (MMP9□), and YFP-positive tumor cells, respectively. Scale bars: 1 mm.

MALDI-MSI and immunofluorescence were thus acquired from different sections of tumors arising in the same mouse model; the observed heterogeneity in marker expression and morphology is consistent with the spatial variation in nucleoside signals revealed by MSI. (file type, PDF)

**Table S1. UHPLC–MS/MS acquisition parameters for nucleoside analysis.**

Multiple reaction monitoring (MRM) transitions (precursor/product m/z), ionization polarity, collision energy, and dwell time used for UHPLC–MS/MS detection of nucleosides. (file type, xlsx)

## AUTHOR INFORMATION

### Author Contributions

SK conceived the study, SK, OK and SYG supervised the study, JW performed experiments with LC-MS and MALDI MS, analyzed the data and wrote the manuscript. SK, OK, SYG, JW conceptualized and wrote manuscript. SR performed IHC stainings, GM developed the LC-MS nucleoside method, analyzed data. NR, KL and FK provided input and material for the study. JW, GN, OK, MG improved the MALDI MS analysis acquired and analyzed data.

## ACKNOWLEDGMENT

The authors of this work want to cordially thank Sylwia Handzik for her excellent technical assistance with tissue preparation and measurement. We are gratefully to Gina Dörpholz and Pia-Katharina Larsen for their support in coordinating activities within the MSTARS project, and to Liliana Mochmann for her assistance in obtaining and coordinating access to clinical FFPE tissue samples and Markus Landthaler for providing standard substances.

## ABBREVIATIONS

FFPE: formalin-fixed, paraffin-embedded
UHPLC-MS: ultra-high-performance liquid chromatography mass spectrometry
MALDI: matrix-assisted laser desorption/ionization mass spectrometry imaging
HESI: heated electrospray ionization
SRM: selected reaction monitoring
ITO: indium-tin-oxide slides
H&E: hematoxylin and eosin
XICs: extracted ion chromatograms
TIC: total ion count
MeOH: methanol
dAde: deoxyadenosine
ROC: receiver operating characteristic
AUC: area under the curve
5-Me_Cyt\2O-Methyl_Cyt: 5-methylcytidine\2O-methycytidine
TNBC: triple negative breast cancer
CHCA: α-cyano-4-hydroxycinnamic acid
LCM: laser capture microdissection.

