## Supplementary figures and images for "Integrating UHPLC-MS and MALDI-MSI for Spatial Nucleoside Profiling in FFPE Breast Cancer: A Multimodal Molecular Pathology Framework"

### SFigure1

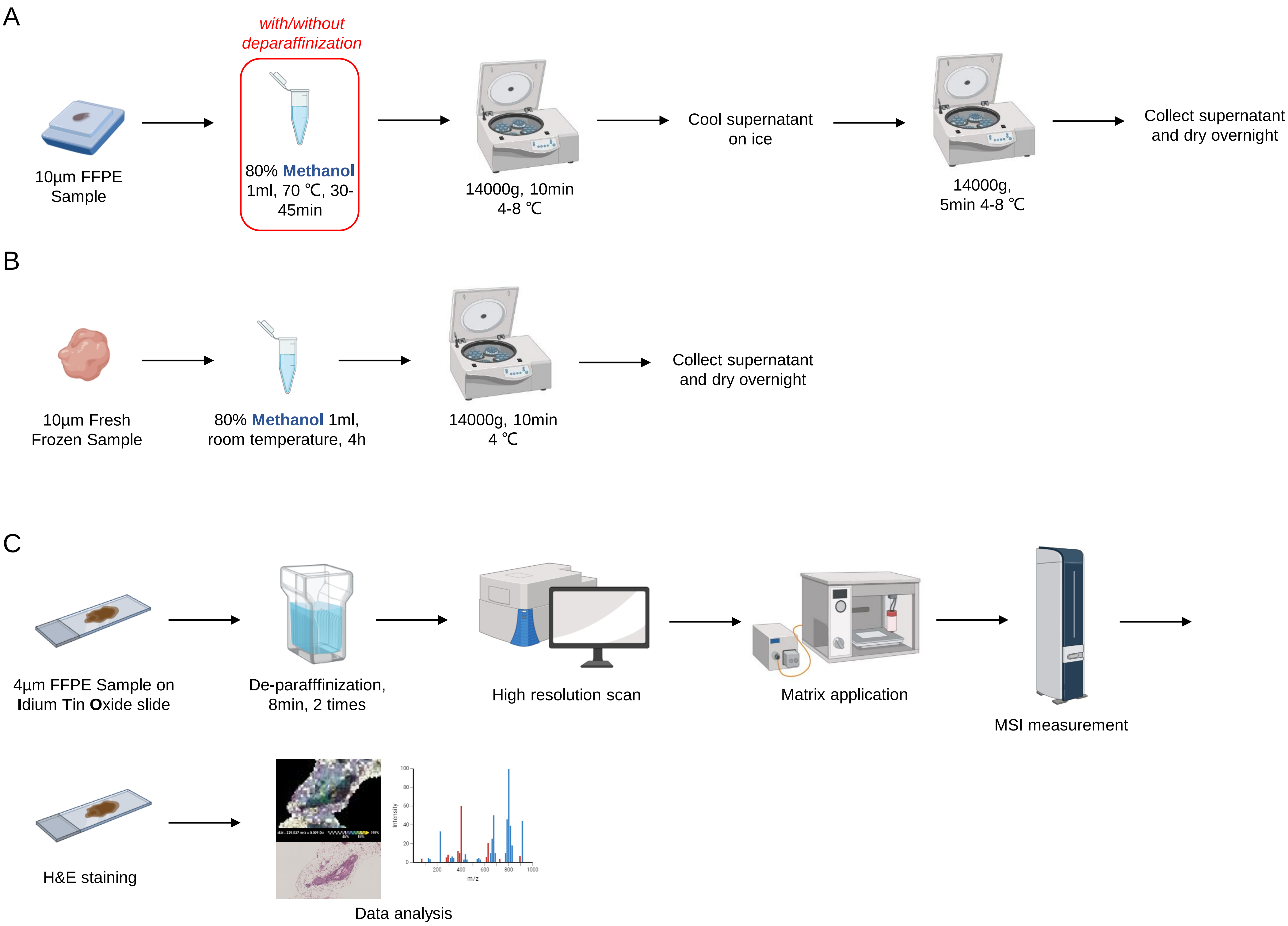

### SFigure2

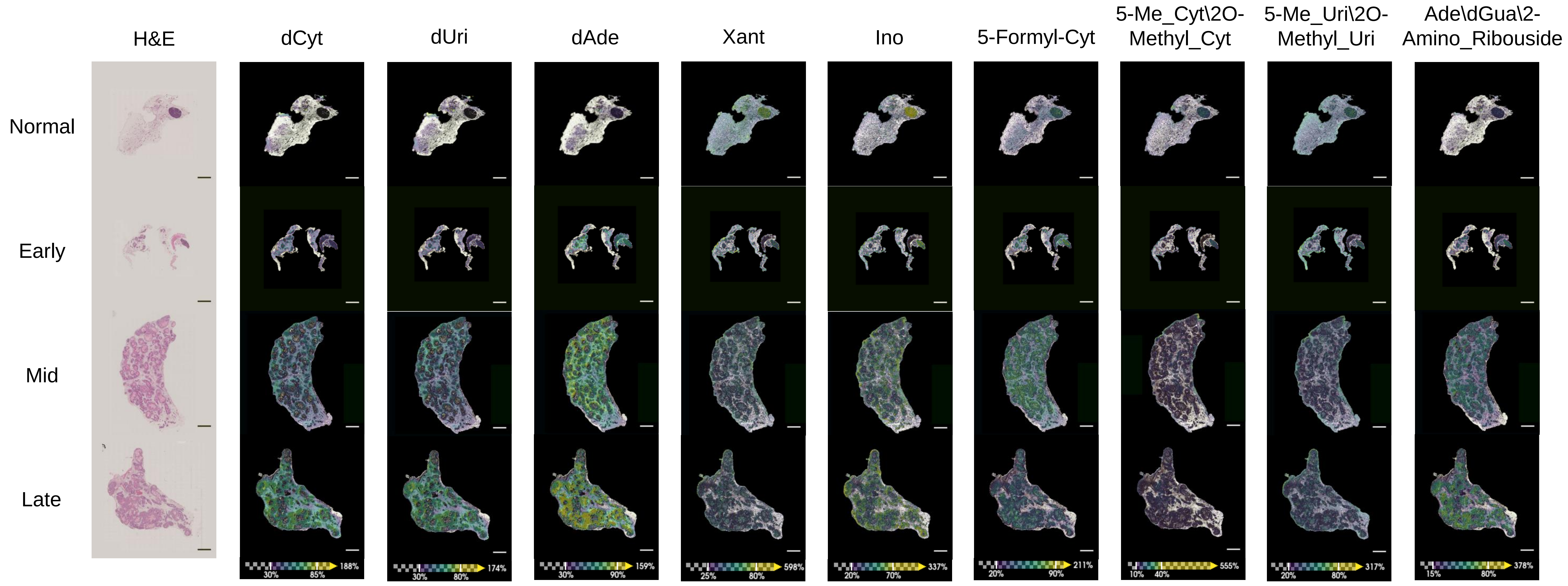

### sFigure3

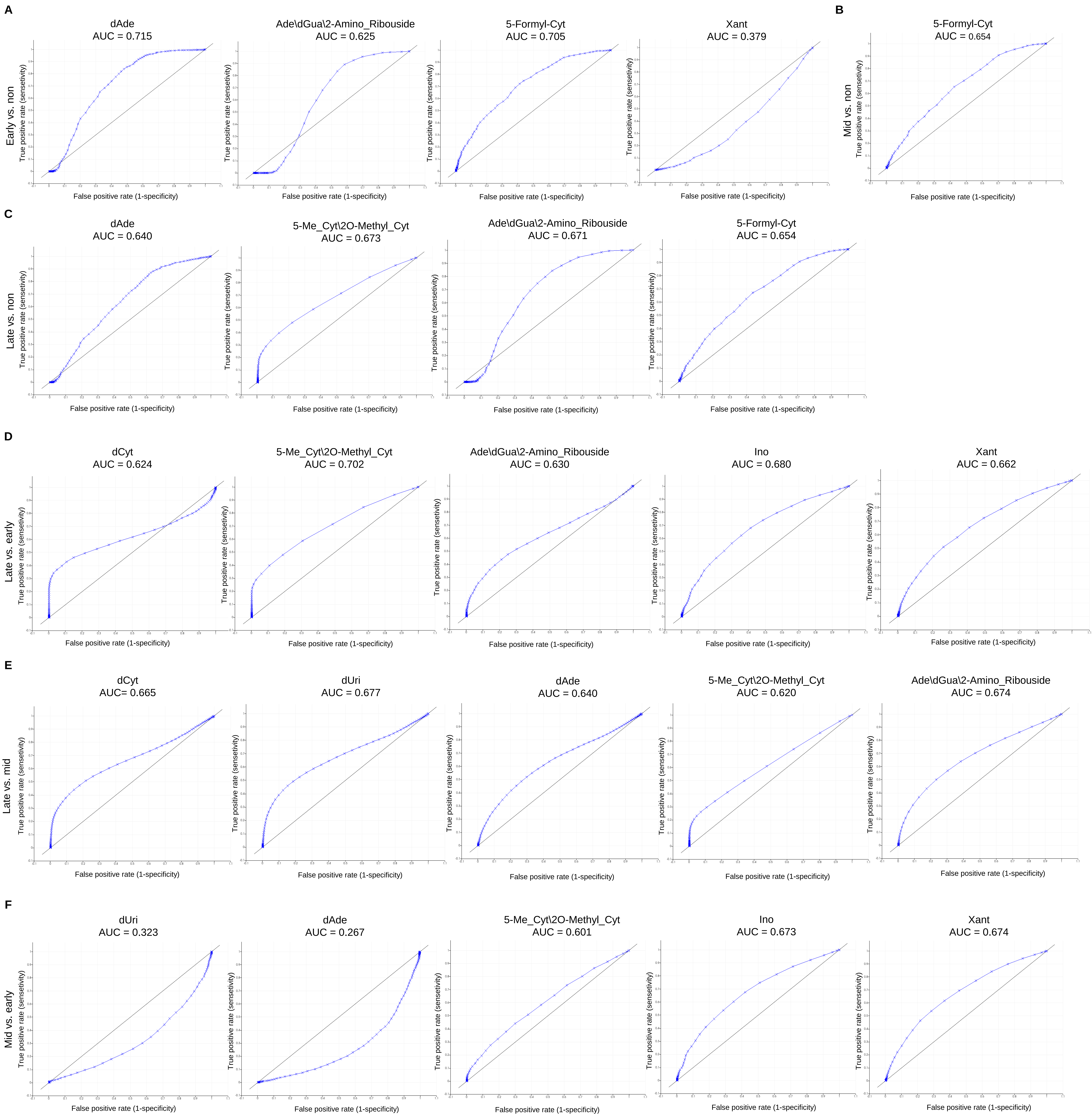

### SFigure4

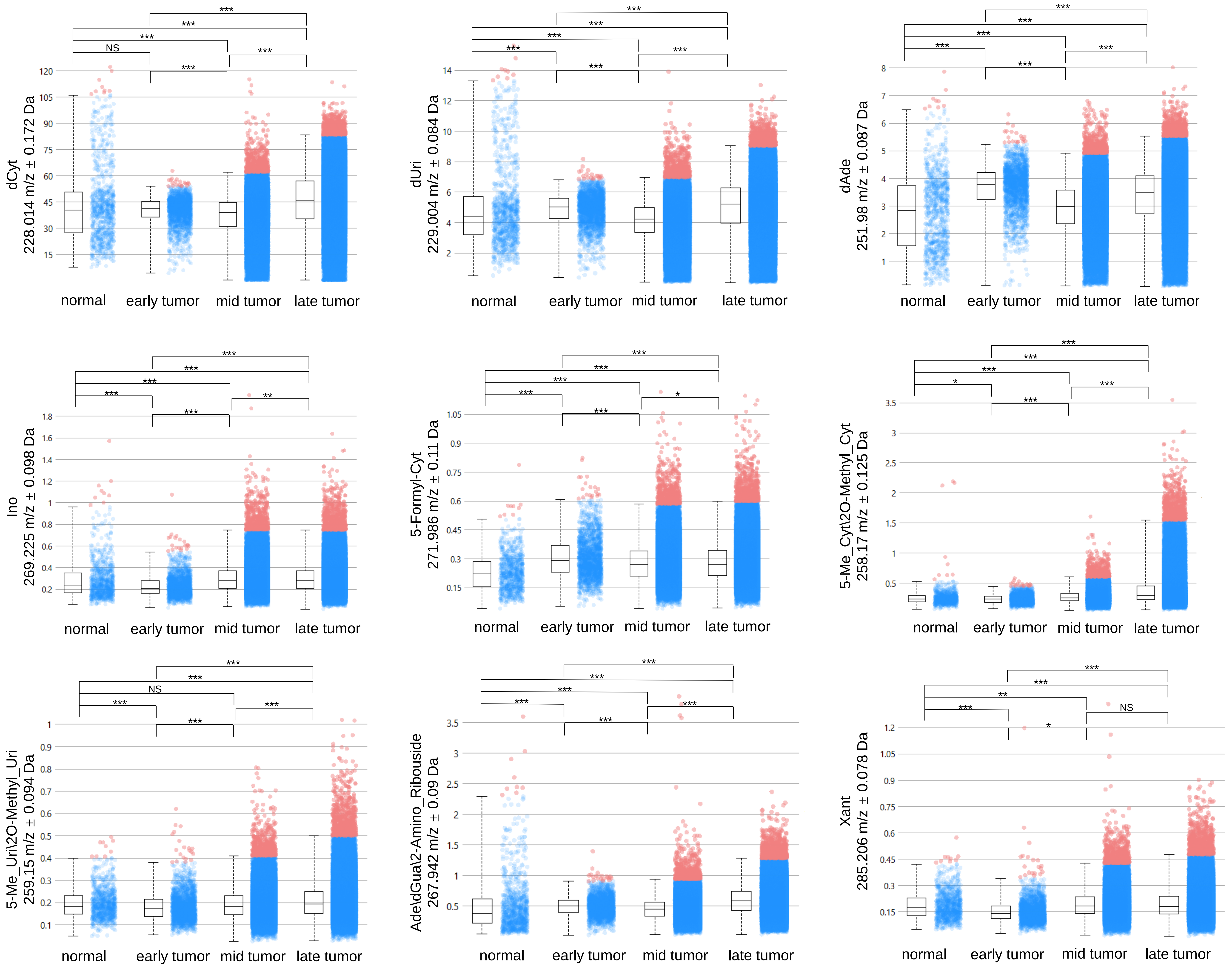

### SFigure5

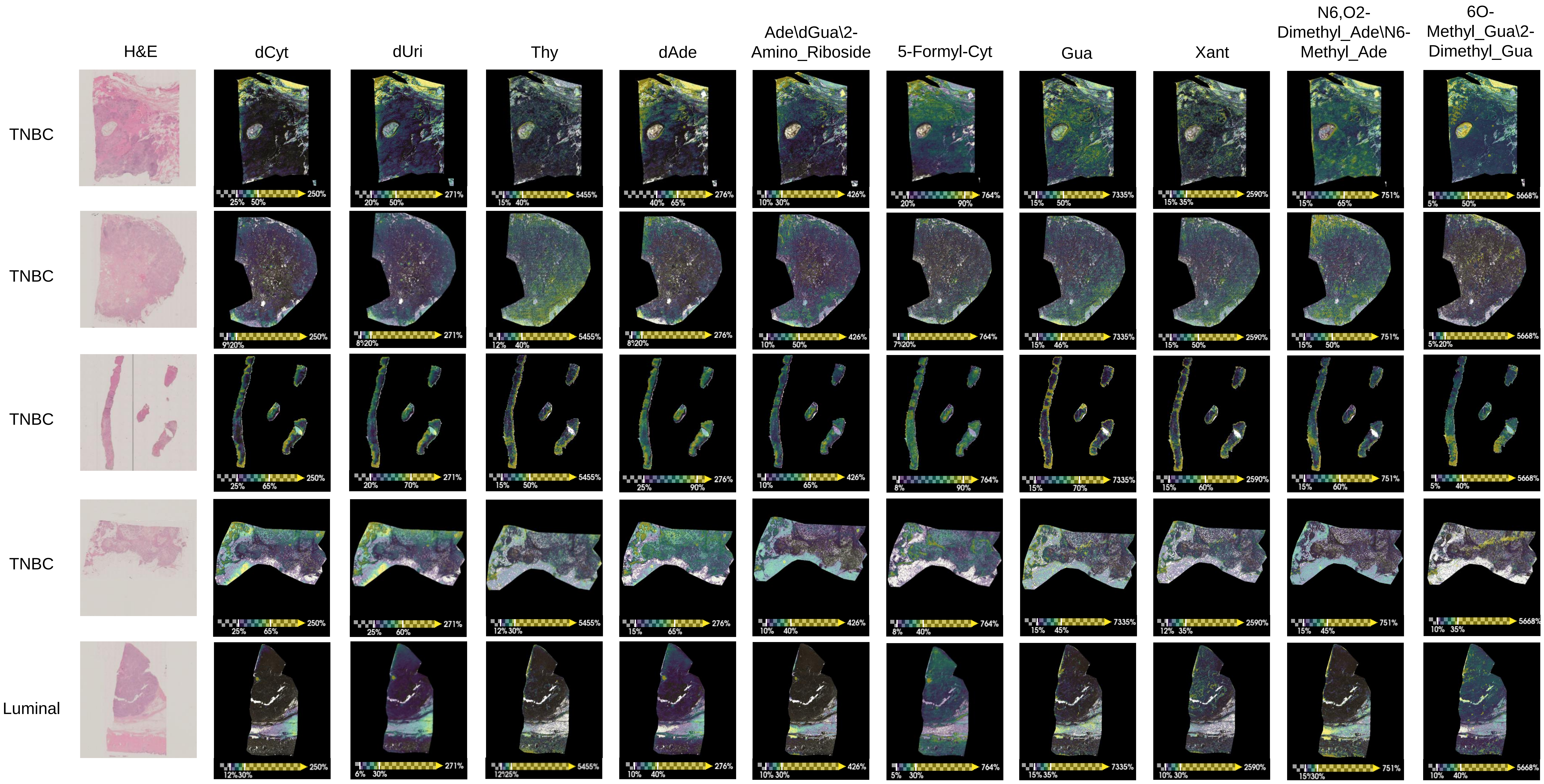

### SFigure6

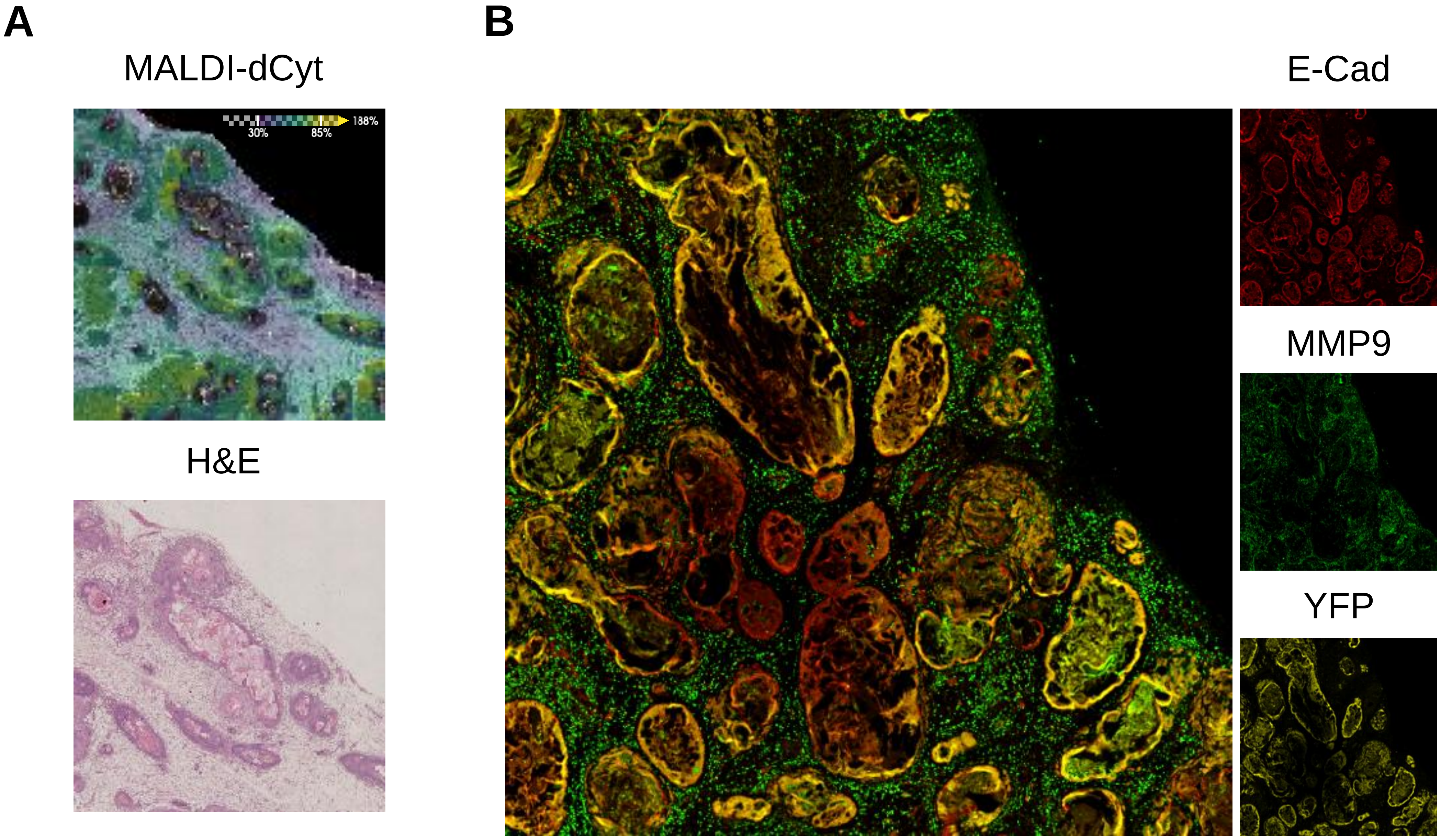
